# Dorsomedial prefrontal neural ensembles reflect changes in task utility that culminate in task quitting

**DOI:** 10.1101/2020.11.05.370536

**Authors:** Blake S. Porter, Kristin L. Hillman

**Author notes:** **Corresponding Author:** Blake S. Porter.

## Abstract

When performing a physically demanding behavior, sometimes the optimal choice is to quit the behavior rather than persist in order to minimize energy expenditure for the benefits gained. The dorsomedial prefrontal cortex (dmPFC), consisting of the anterior cingulate cortex and secondary motor area, likely contributes towards such utility assessments. Here, we examined how male rat dmPFC single unit and ensemble level activity corresponded to changes in task utility and quitting in an effortful weight lifting task. Rats carried out two task paradigms: one that became progressively more physically demanding over time and a second fixed effort version. Rats could quit the task at any time. Dorsomedial PFC neurons were highly responsive to each behavioral stage of the task, consisting of rope pulling, reward retrieval, and reward area leaving. Activity was highest early in sessions, commensurate with the highest relative task utility, then decreased until the point of quitting. Neural ensembles consistently represented the sequential behavioral phases of the task. However, these representations were modified over time and became more distinct over the course of the session. These results suggest that dmPFC neurons represent behavioral states that are dynamically modified as behaviors lose their utility, culminating in task quitting.

**New & Noteworthy:** When carrying out a physically demanding task animals must continually assess whether to persist or quit. In this study we recorded neurons in the dorsomedial prefrontal cortex (dmPFC) of rats as they carried out a challenging weightlifting task, up to the point of quitting. We demonstrate that dmPFC neurons form a representation of the task that is modified, via a decrease in firing rate, by the decreasing utility of the task that may signal quitting.

## Introduction

When engaged in a certain behavior, animals must continuously assess whether to persist in that behavior or change course. During foraging, for example, animals face recurring decisions as to whether to stay in their current patch of diminishing food or to incur the costs of exploration to search for a more abundant patch (1, 2). A third option also exists: rather than adjusting the current course of action, an animal could quit the behavior all together. The decision to quit may be driven by a multitude of factors including physiological states (e.g., satiation, fatigue), the cognitive assessment of diminishing utility, and inter-individual trait variability. However, the neurophysiological processes that underlie quitting behaviors are not well understood.

The prefrontal cortex is likely a critical region in quitting processes due to its role in encoding action-outcome contingencies and using these associations to drive goal-directed behaviors (3–6). In particular, the anterior cingulate cortex (ACC or Anterior Cg1 in rodents) plays a specialized role in evaluating the physical effort costs associated with certain behaviors; this enables net utility determinations and behavioral motivation towards difficult but worthwhile goals (7–10). The ACC is well connected to sensorimotor areas which may translate decisions into actions (11–14).

The secondary motor area (M2) is laterally adjacent to Cg1 in rodents and is considered a higher order motor area that participates in action plan selection based on utility (9, 15–18). Together, these two regions in the rat dorsomedial prefrontal cortex (dmPFC) – the anterior Cg1 and M2 – demonstrate important roles in determining the utility and selection of behaviors that maximize benefit to the animal. However, it is not well-understood how neurons in these areas respond to a prolonged task where quitting eventually becomes the most beneficial action. In such a scenario, the continued time and energy costs to pursue the challenging behavior are no longer worth the payout (e.g., calories gained).

In order to investigate quitting behaviors we recently developed an effortful weight lifting task (WLT) for use with laboratory rats (19). To date, previous electrophysiological studies on cost-benefit decision making have usually not been structured to allow a subject to quit outright or, if subjects do quit, the quitting behavior may not be studied in-depth. Generally, subjects are provided with different cost-benefit actions (e.g., operant actions involving variable effort in rodent studies, (20–22) or cost-benefit contingencies (e.g., stimulus-outcome pictorials in primate studies, (23)) to choose between, and subjects make recursive decisions over a fixed number of trials or a fixed duration. If subjects do not carry out the requisite number of trials or duration, these data may not be analyzed. In our WLT, rats are challenged to pull a weight-adjustable rope a fixed distance in order to earn rewards. Weight-adjustment allows for dynamic control over the effort associated with the rope-pulling behavior. Furthermore, the animals were free to quit the task at any point.

Here we recorded single units from the dmPFC while rats carried out the WLT in order to characterize how neural activity patterns relate to quitting behaviors. Two different paradigms of the WLT were used: a progressive weight (PW) version where the weight increased over time, and a fixed weight (FW) version that used a constant, moderate weight. In the PW version, task utility declines due to the increasing costs (weight) while task utility in the FW version declines due to the internal states of the animal (e.g., fatigue, satiation). We predicted that dmPFC neurons would be responsive to our effort-based task in that different neurons should represent the actions, outcomes, and salient behavior transitions associated with performing the task. Furthermore, these representations should be modulated by changes in utility, brought about by heavier weight-adjustments and/or fatigue. We predicted that Cg1 neurons would be more responsive during reward phases to monitor action-outcome contingencies, while M2 neurons would be more responsive to rope pulling and associated effort. Lastly, we predicted that neural populations in the dmPFC would undergo state transitions that reflect the rats’ behavioral transitions towards the point of quitting.

## Materials and Methods

### Subjects

Male Sprague-Dawley rats (n=6, weighing between 400 - 480 g at the time of experiment; Hercus Taieri Resource Unit, New Zealand) were single housed in individually ventilated cages measuring 38 x 30 x 35 cm (Tecniplast, Italy). Animals were maintained on a 12 h reverse light cycle (07:00 to 19:00), with experimentation occurring during the animals’ dark, active phase. Water was available *ad libitum* throughout training and experimentation. Daily chow (Teklad diet; Envigo, USA) was rationed to promote food motivation for training and during experimentation. Rats were weighed twice per week and daily food rations were adjusted to maintain each animal’s body weight ≥ 85% of their free-feeding body weight. All procedures were approved by the University of Otago’s Animal Ethics Committee, protocol 91/17.

### Training

Prior to surgery, animals were trained in the WLT as previously described (19). Briefly, the WLT is performed in a 120 x 90 x 60 cm arena (Figure 1a). An animal is progressively shaped to pull a rope 30 cm to trigger the dispensing of 0.25 mL of 20% sucrose solution. The rope can be weighted from 0 to 225 g to increase the effort demands of the task, however only 0 g and 45 g are used in the training phase. Once an animal can perform 20 successful pulls (10 pulls on 0 g followed immediately by 10 pulls on 45 g) in under 10 minutes, they are considered trained in the WLT and ready for surgery. Rats took an average of 26.3 days (Standard Deviation (S.D.) = 0.7 days) to reach the 20 successful trial criterion.

**Figure 1:**
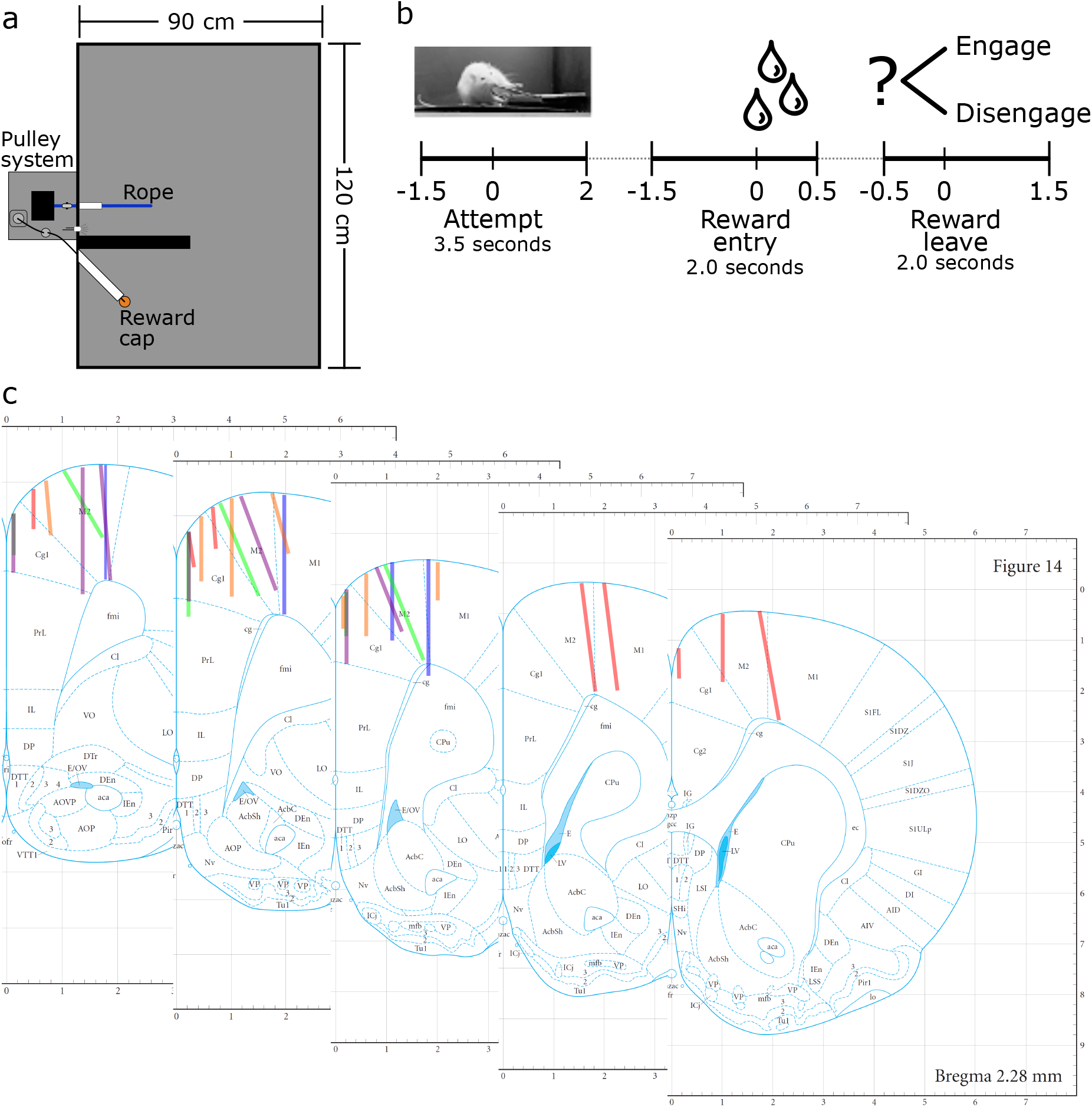
Task overview and histology. a) Weight lifting task schematic as reported in Porter and Hillman (2020). b) Trial breakdown illustrating the three phases of interest. In the Attempt phase, rats are physically pulling the rope. In Reward entry, rats enter the reward cap area and consume sucrose reward if present. In the Reward leave, rats leave the reward cap area and choose to re-engage in a rope pull or disengage. c) Histological reconstruction of tetrode positions. Each color represents the tetrodes from one rat. Tetrodes spanned +3.72 mm to +2.28 mm relative to Bregma. Schematics modified from Paxinos & Watson (2007).

### Microdrives

All rats were implanted with Halo-10 Microdrives (Neuralynx, USA) consisting of eight independently drivable tetrodes. Tetrodes were made from 18 micron Nichrome Formvar insulated wire (A-M Systems, Inc., USA). Tetrodes were gold plated over three to five days using a nanoZ (White Matter LLC, USA) such that electrode impedances were 200-300 kOhms prior to surgery. Tetrodes were secured to Neuralynx EIB-36’s via gold pins to interface with our headstage (see *Data collection*, below). Neuralynx Standard A exit tips were used and tetrodes were positioned in the exit tip to minimize medial-lateral spread when implanted in the brain.

### Surgery

Anesthesia was induced with 5% Isoflurane in oxygen and then maintained at 2-3% throughout the surgery. Rats were mounted in a Kopf (David Kopf Instruments, USA) stereotaxic frame with non-puncture ear bars. One mL of 2% Lidocaine diluted in 4 mL of 0.9% saline was injected subcutaneously above the skull. A midline incision was made and the skin over the skull retracted. A 3 mm diameter craniotomy was made centered at + 2.5 mm anterior-posterior and ± 0.7 mm medial-lateral from Bregma (Paxinos and Watson, 2007). Five rats were implanted on the right hemisphere and one on the left hemisphere. The dura was removed and the microdrive was lowered onto the surface of the brain via a stereotaxic arm such that the most medial tetrode was next to the central venous sinus. Two 4 mm long, 1 mm wide stainless steel screws, bilaterally placed over the cerebellum, served as grounds. Six to eight additional 3 mm long, 1mm wide stainless steel screws were then put partially into the skull and the implant was secured to the skull and skull screws with ultraviolet cure self-etching dental cement (Maxcem Elite™, Kerr Corp., USA). Sutures were used to close the skin around the implants.

Post-operatively (post-op), rats were subcutaneously administered prophylactic antibiotic (30mg/kg of Amphoprim) and an anti-inflammatory (5 mg/kg of Carprofen) and recovered in a heated cage until they regained consciousness and were active. Rats were then returned to a clean home cage to recover for 10 days. The anti-inflammatory was administered for a minimum of two additional days post-op and as needed if rats showed any signs of pain. After the 10 days of post-op recovery, rats were re-introduced to the WLT and tested to criterion to ensure that they could still carry out the task and habituate to their implants. Rats took an average of 9.2 (S.D. = 4.9) days to return to criterion. Once rats reestablished criterion performance, experimentation started.

### Data collection

Single unit data was collected using a Neuralynx Digital Lynx SX system with Cheetah software (version 6.3.2) via a Neuralynx HS-36-LED pre-amplifier headstage connected to a Neuralynx Saturn commutator. Data from the 32 electrodes were sampled at 32 kHz and bandpass filtered between 600 and 6,000 Hz. Every tetrode was referenced to an electrode on a different tetrode with no spiking activity. Single unit action potentials were detected online with a threshold of 65 µV and stored for offline spike sorting. Spikes were sorted offline using Plexon (USA) Offline Sorter (version 4) by primarily relying on peak-valley amplitude and principal component analysis. Only well-isolated clusters of waveforms, likely to be from the same neurons, were included for analyses. Tetrodes were not moved day-to-day if they had single units on them. If a tetrode did not have any single units it was advanced approximately 32 microns at the end of a recording session.

Each headstage contained two LEDs which enabled the animal’s position to be tracked by Neuralynx’s Cheetah software via an overhead camera (JAI Inc., USA). Video was sampled at 25 frames per second. An Arduino Uno (www.arduino.cc) was used to detect rope pulls and to dispense sucrose solution reward; TTL signals of all events were sent to the Digital Lynx SX system for timestamping and storage.

### Progressive weight paradigm

Rats completed two WLT paradigms: a progressive weight (PW) paradigm and a fixed weight (FW) paradigm (19). In the PW paradigm, a session started with a two minute pre-baseline period where the rat was placed in the WLT arena but the rope was not extended into the environment. After the pre-baseline, the rope was extended with 0 g of weight attached and the rat could begin earning rewards; each successful 30 cm pull dispensed 0.2 mL of 20% sucrose solution. After every 10 successful (rewarded) pulls of the rope, 45 g of weight was added to the rope until the rat quit or 225 g was reached. If a rat completed 10 trials of 225 g, a clip was placed on the rope in order to prevent the rat from pulling the weight high enough to trigger reward dispensing. However, no rats in this study completed 10 trials of 225 g.

Quitting was defined as the rat making no attempts to pull the rope for two full minutes. When a rat quit the rope was retracted from the apparatus at the end of the two minute ‘quit period’, and a two minute post-baseline period started. Lastly, after the post-baseline period, two sucrose rewards were manually dispensed to check if the rats were satiated or not. Rats had one minute to consume the sucrose reward once dispensed. Every rat on every session consumed these manually dispensed rewards, suggesting that satiation was not occurring. After the sucrose satiation check the recording session was stopped and the animal was returned to its home cage. Rats ran one session per day.

### Fixed weight paradigm

In the FW paradigm, a session again started with a two minute pre-baseline period, followed by the rope being extended into the arena with 0 g of weight attached. After 10 successful (rewarded) pulls on 0 g, the rope weight was immediately switched to a fixed weight, calibrated to each specific rat. The initial fixed weight was the maximum weight that the rat had completed in the PW paradigm (i.e., the weight on which they had successfully performed 10 rewarded pulls). Rats were allowed to complete as many trials as they wanted on the FW until they quit or until 60 minutes elapsed, whichever came first. After 2-3 days (average 2.7 days, S.D. = 0.4 days), the weight was adjusted to be 45 g heavier or lighter depending on the rats’ performance: if rats were working for fewer than 15 minutes their weight was reduced whereas if rats were working for more than 15 minutes their weight was increased. Rats then carried out the adjusted FW paradigm for 2-5 further days (average 3.2 days, S.D. = 1.0 days) depending on their performance and the number of single units being recorded.

### Overall analyses

All analyses were carried out using Matlab (version R2018a; The MathWorks Inc., USA). Colorbrewer (https://github.com/axismaps/colorbrewer/) and mpl colormaps (https://bids.github.io/colormap/) were used for graphical color maps.

### Behavioral analyses

Behavioral metrics were calculated as previously reported (19). Briefly, for the PW paradigm, the quit weight was calculated as the highest weight in a given session on which the rat made at least one rope pulling attempt. For the FW paradigm, the quit duration was calculated as the time from when the FW was attached (exclusive of 0 g) until the last pull attempt.

For both the PW and FW paradigms, quartiles were determined based on the duration of each individual session. For the PW paradigm, the duration was calculated starting from when the 0 g weight was available until the rat quit. For the FW paradigm, quartiles were calculated starting from the time the FW was available (thus excluding the initial period of 10 trials on 0 g). For all FW paradigm analyses this initial 0 g period was excluded so as to avoid comparisons between 0 g-containing quartiles and FW-containing quartiles Thus, for behavioral and neural analyses of the FW paradigm, every quartile has the same weight attached to the rope. Every quartile had to contain at least five completed trials; recording sessions that did not meet this minimum trial criterion were excluded from this manuscript. The four quartiles are abbreviated throughout the manuscript as Q1, Q2, Q3, and Q4 to denote the first, second, third, and fourth quartile.

Trials were defined as the sequence of events beginning with a rope pull attempt followed by reward area entry and concluding with reward area leaving. This provided three task phases: the attempt phase, the reward entry phase, and the reward leaving phase (Figure 1b). The attempt phase consisted of 1.5 seconds before and 2.0 seconds after a pull attempt. Pull attempts were timestamped by a TTL code from the Arduino that logged when the weight was lifted. Reward entry and reward leaving were based on the tracking data obtained from LEDs on the headstage. A 20 x 20 cm area was manually placed around the reward cap for each recording session based on the rat’s tracking data and known cap location. Reward entry was centered on when the rat entered the reward zone and their instantaneous speed dropped below 15 cm/s. The reward entry phase consisted of 1.5 seconds before and 0.5 seconds after reward zone entry. Reward leaving consisted of 0.5 seconds before and 1.5 seconds after the point where the rat left the reward zone. Task-phase durations were chosen based on the distribution of durations across all recording sessions in order to maximize duration but minimize overlap between phases.

Trials were deemed “successful” when the rat pulled the weight sufficiently high enough (30 cm) to trigger a reward, or deemed “failures” when the rat did not pull the weight high enough but still went and sampled the reward cap. Events when the rat pulled the rope but did not check the reward cap were excluded. Trials were also parsed according to whether or not the rat “engaged” or “disengaged” from the task on the subsequent trial. Engagement was determined based on how long it took from the rat leaving the reward cap until the next pull attempt. If the rat made another attempt within 12 seconds the trial was considered an engage trial; anything longer than 12 seconds was considered a disengage trial. Twelve seconds was chosen as a cut off as 75% of all next-attempts occurred within 12 seconds of leaving the reward cap.

The on-task ratio was calculated as the total time a rat spent working at the task divided by the total time of the session when the rope was available. Rats were considered working if they were both within the “work zone” and making attempts to pull the rope. The work zone consisted of a 60 x 64 cm area encompassing the rope pulling area and the reward cap. The reward interval was calculated as the time between earned rewards based on the reward TTL signals. The success-to-attempt ratio was calculated as the number of successful pull attempts divided by the total number of attempts made. For the on-task ratio and success-to-attempt ratio we used a repeated-measures one-way ANOVA to test for significant changes over the quartiles and Tukey-Kramer’s test for post-hoc pairwise comparisons. Due to some rats having no rewards in Q4, and thus no reward intervals, we instead used a Kruskal-Wallis test to measure if reward interval times were affected by quartile and Tukey-Kramer’s test for post-hoc comparisons.

For every trial we calculated the animal’s average speed as they traversed from the attempt zone to the reward zone after an attempt was made. We carried out a one-way ANOVA to determine if there were significant changes in speed over quartiles.

### Neural analyses – single neurons

For every recording session, sorted spikes from Plexon Offline sorter were imported into Matlab with the accompanying tracking and TTL data from that recording session. Neurons were assigned a brain area based on what tetrode they were recorded from. We tested whether or not neurons were task-responsive by randomly sampling every neuron’s firing rate across 10,000 two second epochs drawn from the entire recording session (including the pre- and post-baselines). Neurons were considered to be task-responsive if, during at least one task-phase, their firing rate was significantly different (Wilcoxon rank sum test) to the random sample and they had a firing rate of at least 0.5 Hz. Furthermore, to avoid including neurons that may have been lost or gained over the course of a recording session, a neuron also had to have a non-zero firing rate in at least one task-phase for all four quartiles. Neurons that did not meet this criteria were not included for analysis. Spike raster plots and peri-event time histograms were created for task-responsive neurons for each of the three task-phases across all trials. Spike times were aligned to the center of the task-phase event. Histograms were created with 100 ms bins across each task-phase. For each task-phase, a Chi-squared test against an equal distribution was used to test if there was an unequal distribution of task-responsive neurons across brain areas.

Neurons were determined to be phasic or tonic based on how many bins of a trial they fired an average of 2 standard deviations above their overall mean firing rate. Neurons that fired for more than 2 bins but less than 10 bins were deemed phasic while all others were deemed tonic. Neurons were considered to have an inhibitory response to the task phases if they had an average response of less than −0.5 standard deviations for 10 or more bins. Neurons that did not fit this criteria were deemed excitatory.

### Neural analyses – population analyses

All task-responsive neurons from all sessions were pooled together for further analysis. A sequence plot was created with all neurons where each neuron’s binned trial firing rates were z-scored then averaged across all trials in each 100 ms bin. All neurons were then sorted based on the trial bin that contained their maximum firing rate. For graphical purposes only, every neuron’s activity was normalized to be between 0 and 1 where 1 corresponded to their maximum z-scored firing rate and 0 corresponded to their minimum z-scored firing rate. All analysis was done on the z-scored firing rates. A cross-correlation matrix was created using the sequence plot where the Pearson’s correlation was calculated pairwise for every 100 ms bin of a trial.

Principal component analysis (PCA) was carried out on the z-score averaged firing rates of all task-responsive neuron in 100 ms bins across the trial using Matlab’s pca function. Trials were the same as described above, consisting of the 3.5 second attempt phase, 2.5 second reward entry, and 2.5 second reward leave phase for consistent behaviors and durations. Thus PCA was carried out once on a matrix where every column was a neuron and every row was the neuron’s average z-scored firing rate for every 100 ms bin of a trial. PCA was also carried out on the successful versus failed trials, engage versus disengage trials, and quartiles. In these cases, each task-responsive neuron’s firing rates were binned into 100 ms bins, z-scored across all trial types, and then all neurons were pooled together. A matrix was then constructed where every column was a neuron and every row was a 100 ms bin of a trial for each trial type (successful and failed, engage and disengage, and each of the quartiles). PCA was ran once on the matrix resulting in the same principal component space for all trial types. We computed the distance between the PCA score trajectories by calculating the distance between each 100 ms bin using the first three principal components. As a control, we also randomly split trials of a recording session into halves (for engage versus disengage comparisons, and success versus fail comparisons) or quarters (for quartile analyses) and averaged a neuron’s activity over the random trials. This was carried out 1,000 times per session. For every iteration, all neurons were pooled together, PCA carried out, and the distance between each 100 ms bin calculated. For the comparisons of engage versus disengage and success versus fail, a paired t-test was used to determine if the observed PCA distances were significantly different from the mean of the 1,000 random PCA score distances. A one-way ANOVA, with Tukey-Kramer’s post-hoc, was used to test whether or not the observed PCA score distances across the three task-phases were significantly different.

For graphical purposes only, the PCA score plots were smoothed with a 500 ms moving average within each task-phase when comparing multiple trajectories. The PCA trajectories of all trials from the PW and FW paradigms (Figures 4c and 7c) are plotted with the raw scores. All distance calculations and statistics were done on the raw PCA scores.

Kruskal-Wallis tests were used on each task-responsive neuron to test firing rate variability across quartiles for each task-phase. To keep trial numbers consistent across sessions, we limited analysis to five trials in each quartile. For Q1, the first five trials were used. For Q2 and Q3, the middle five trials were used. For Q4, the last five trials were used. We then pooled the quartile averaged z-scored firing rates of all neurons with a significant (p < 0.05) effect for quartile for each task-phase. For each task-phase we determined what quartile each neuron was most active in based on their averaged fire rates in each quartile. We carried out a Chi-squared test on the observed distribution of peak activity across the four quartiles versus and equal distribution across all quartiles. Repeated-measures one-way ANOVA with a post-hoc Tukey-Kramer’s test was used on the populations of neurons for each task-phase, with quartile as a factor.

As a control, we iteratively subsampled proportions of the total population of neurons and repeated the PCA analyses. We carried this out 1,000 times for each subsampled proportion and averaged the iterations together. The average PCA trajectories were visually inspected and their pairwise distances were tested against distances from the above described randomly sampled PCA distances.

Neuronal data was also broken up into two minute epochs across the entire recording session, including the pre- and post-baseline epochs where the rope was not available to the rat. Four more epochs consisted of the four quartiles. For quartile one, the first two minutes of the 0 g (PW paradigm) or the fixed weight (FW paradigm) was used. For quartiles two and three, the middle two minutes of the quartile were used. For quartile four, the two minutes preceding the rats’ final attempt were used. The quit epoch consisted of the two minutes proceeding the rats’ final attempt where the rope was still available but the rat was not making attempts. For each task-responsive neuron a z-scored firing rate was calculated for each two minute epoch and all neurons were pooled. Pairwise cross-correlations were calculated using Pearson’s correlation. Matlab’s pdist function was used to calculate the pairwise correlation distances between epochs. Matlab’s linkage function was then used to hierarchically cluster the correlation distances based on their average correlations. Matlab’s dendrogram function was used to plot the clustering.

As an additional control analyses, we tested whether session duration had an impact on the changes in neural activity that we observed across quartiles. First, we tested if the magnitude of change seen in neuron firing rates was influenced by time. If dmPFC neural activity was changing due to time (24), we would expect a greater change in activity for longer recording sessions. We calculated the difference in firing rate for each neuron from Q1 to Q4 for each task-phase and ran a linear regression with session duration as the explanatory variable. We did this for all task-responsive neurons and then again for only task-responsive neurons that had a significant effect of quartile.

We carried out a second control analysis for time where we again found the difference between the mean firing rate of Q1 to Q4 in the 2 minute epoch periods. We again fit a simple linear regression with time as the predictor.

### Histology

After completion of all behavioral paradigms, rats were deeply anesthetized with Isoflurane and transcardially perfused with 4% paraformaldehyde (PFA) in phosphate-buffered saline (PBS). All tetrodes were wound up before the brain was removed to avoid damage. Brains were then cryoprotected in 30% sucrose in 4% PFA PBS until they sunk. Brains were sectioned at 40 µm using a Cryostat (CM1950, Lecia Biosystems, LLC., Germany). Brain sections were then mounted, stained with Thionine, and imaged with a light microscope (Lecia Biosystems, LLC.). Tetrodes were mapped based on their tracks in reference to (25) brain atlas, microdrive turn logs, and their relative locations in the microdrive (Figure 1c).

## Results

### Progressive weight behavior

Rats first carried out the PW paradigm; here the rope weight started out at 0 g and was increased by 45 g every 10 successful pulls, to a maximum weight of 225 g. Rats were free to quit the task at any point. Six rats performed a total of 30 sessions of the PW task (3-7 sessions per rat; average 5.0 ± 1.7 S.D.). Consistent with our previous studies (19), rats were more likely to quit on heavier weights (Figure 2a). No rat was able to complete 10 successful pulls of 225 g. Due to the inter- and intra-individual variance of quit weights across sessions, each session varied in time duration and number of trials. Sessions varied in length from 874 to 2,056 seconds (median = 1,589 ± 373). Across all sessions rats earned from 25 to 55 rewards (median = 47 ± 8). Because of this variability, we broke each session into quartiles based on time and used those quartiles to analyze behavioral performance.

**Figure 2:**
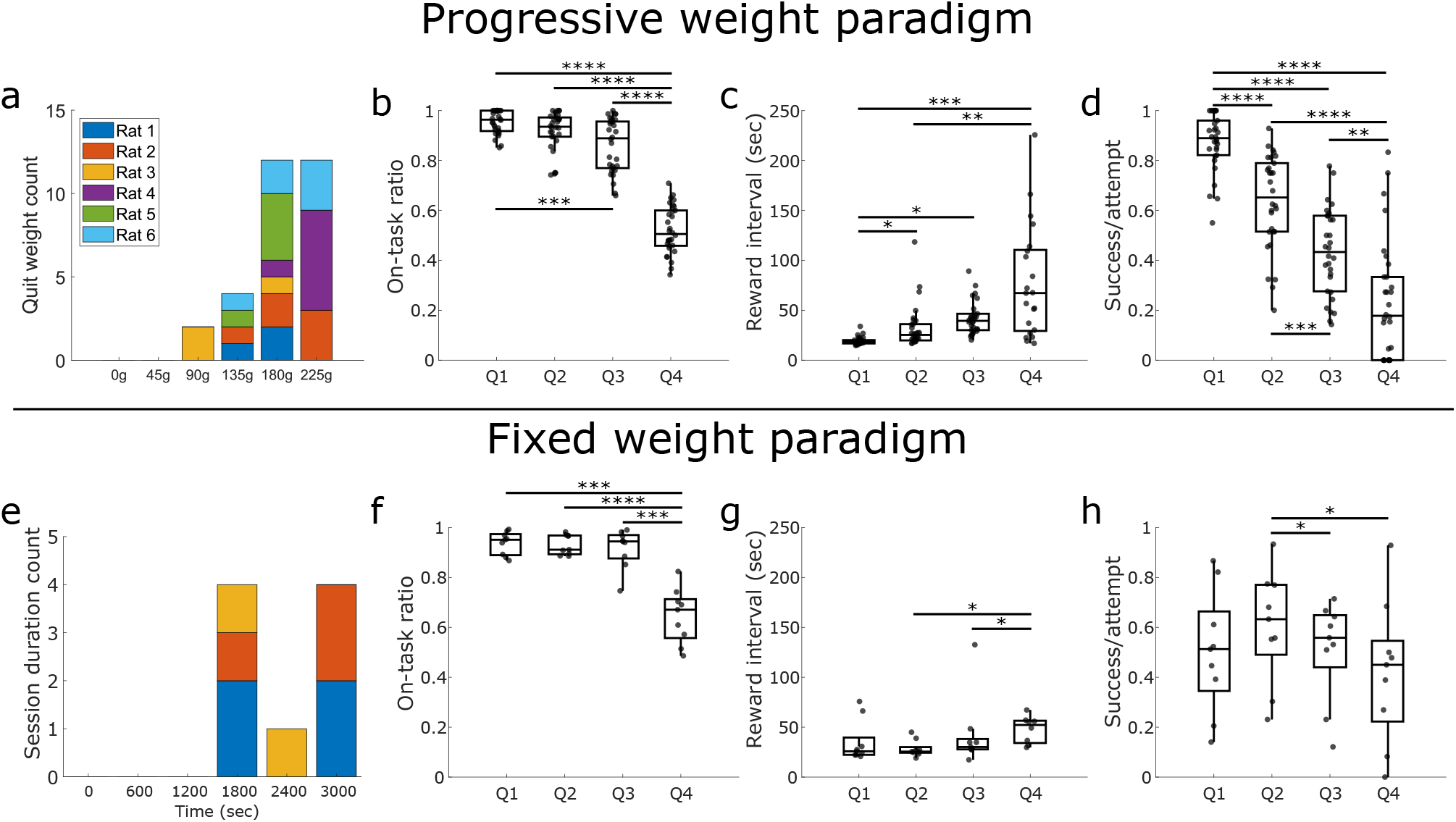
Behavioral measures on the PW and FW paradigms. a-d) Behavioral metrics for the PW task. e-h) Behavioral metrics for the FW task. For both tasks, each recording session was broken up into four equal duration quartiles, Q1-Q4. Histograms illustrate the weight that the rats quit on in the PW (a) or the duration until quitting in the FW (e). Each rat is assigned a different color. The time on-task ratio was calculated as the time rats spent working (making attempts) divided by the total time in each quartile. The success-to-attempt ratio was calculated as the number of successful attempts divided by the total number of attempts for each quartile. Box plots represent the first and third quartile with the middle line representing the median. Error bars represents ± 1.5 times the interquartile range. * p < 0.05, ** p < 0.01, *** p < 0.001, **** p < 0.0001.

We computed a time-on-task ratio that reflected the proportion of time rats spent engaged in the task. A ratio of 1 indicated that the rats were regularly making attempts to pull the rope while a ratio of 0 reflected rats making no attempts. Across quartiles, rats spent significantly less time on-task as the session progressed into the third and fourth quartiles (F (2.6, 76.4) =177.0, p < 0.0001; Figure 2b). When rats were on-task and working, it took longer to earn rewards as the session progressed (H (3, 107) =56.0, p < 0.0001; Figure 2c). Increases in reward interval, notably in Q4, were likely in part due to an increasing number of failed rope pulls as weights got heavier. Across quartiles, there was a significant decrease in the success-to-attempt ratio (F (2.4, 68.4) = 88.1, p < 0.0001; Figure 2d). Rats also slowed down in their reward approach on the fourth quartile relative to the other three quartiles (F (3) = 19, p < 0.0001; Supplemental Figure S1a https://doi.org/10.6084/m9.figshare.14515644).

### Fixed weight behavior

On the FW paradigm, the rope weight started out at 0 g and after 10 successful pulls the rope weight was set to a moderate FW, calibrated to each rat (see Methods). Rats could perform as many weighted rope pulls as desired, up to a maximum duration of 60 minutes. Rats were free to quit the task at any point. Because we set a minimum trial number criterion for data analysis inclusion (see Methods), only nine sessions of the FW task were available for analysis (three rats performed these nine sessions, 2-4 sessions per rat; average 3.0 ± 1). Quit durations ranged from 1,803 to 3,548 seconds (median = 2,711 ± 571 seconds Figure 2e) and rats earned from 24 to 106 rewards (median = 81 ± 27 rewards) on the FW.

Similar to the PW paradigm, rats spent significantly less time on-task as the FW sessions progressed (F (1.7, 13.3) = 47.5, p < 0.0001; all pairwise comparisons to Q4, p’s < 0.0001; Figure 2f). There was a main effect for quartile on reward interval (H (3, 31) = 9.8, p = 0.02; Figure 2g), with Q4 having a higher reward interval than Q3 (p = 0.030) and Q2 (p = 0.035). There was a significant main effect of quartile for the success-to-attempt ratio (F (2.1, 17.0) = 5.9, p = 0.01; Figure 2h). However, post-hoc pairwise analyses only showed modest differences between quartiles: success-to-attempt ratio was significantly higher on Q2 compared to Q3 (p = 0.041) and Q4 (p = 0.043). Compared to behavior in the PW task, where weights incrementally increased by 45 g every 10 trials, rats may have struggled with the FW initially in Q1 where the weight suddenly increased from 0 g to the FW value greater than 135 g. This likely accounted for the lower success rate and greater reward interval observed in Q1 in the FW task. There was no change in the rats’ speed to leave the attempt zone and approach the reward zone across quartiles (F (3) = 0.15, p = 0.93; Supplemental Figure S1b https://doi.org/10.6084/m9.figshare.14515644).

### Neuronal task selectivity – Progressive weight paradigm

For neuronal analysis we focused on three task-phases of interest: the attempt phase, the reward entry phase, and the reward leaving phase (see Methods and Figure 1b). In the PW paradigm, we recorded a total of 360 neurons across the 30 recording sessions. Based on histological reconstruction (see Figure 1c), 123 of these neurons were from the Cg1, 93 were between Cg1-M2, and 144 were from Motor cortices 2/1 (M2/1).

A variety of activity patterns were observed within and across task-phases as well as over time (Figure 3a-h). Some neurons were especially active during the attempt phase (3a, b). The Cg1 neuron shown in Figure 3a appeared to have increased its firing rate prior to the attempt and then fired rhythmically at approximately 2 Hz. The motor cortex neuron in Figure 3b was highly selective to the attempt phase with high firing right before the attempt followed by a rhythmic firing pattern during rope pulling. Other neurons had phasic firing to very specific time points of a trial. For example, the Cg1-M2 neuron shown in Figure 3c fired just prior to reward entry while the M2/1 neuron in Figure 3d fired as the rat left the reward zone. Furthermore, some neurons showed multiple phasic peaks such as the motor cortex neuron in Figure 3e that was active both right before reward entry and at reward leaving. More complex activity profiles were also observed such as the Cg1 neuron shown in Figure 3f that fired tonically during all three phases but appeared to be briefly inhibited prior to reward entry and after reward leaving. Some neurons showed a clear change in firing rate over the course of the recoding. For example, the neuron in Figure 3g was highly active before reward entry on early trials (top lines of firing rate raster plot) but decreased its firing rate at reward entry as the weight became heavier over the session (moving vertically down the raster plot). The motor cortex neuron shown in Figure 3h doesn’t have a clear preference for a trial-phase but does increase in firing over time.

**Figure 3:**
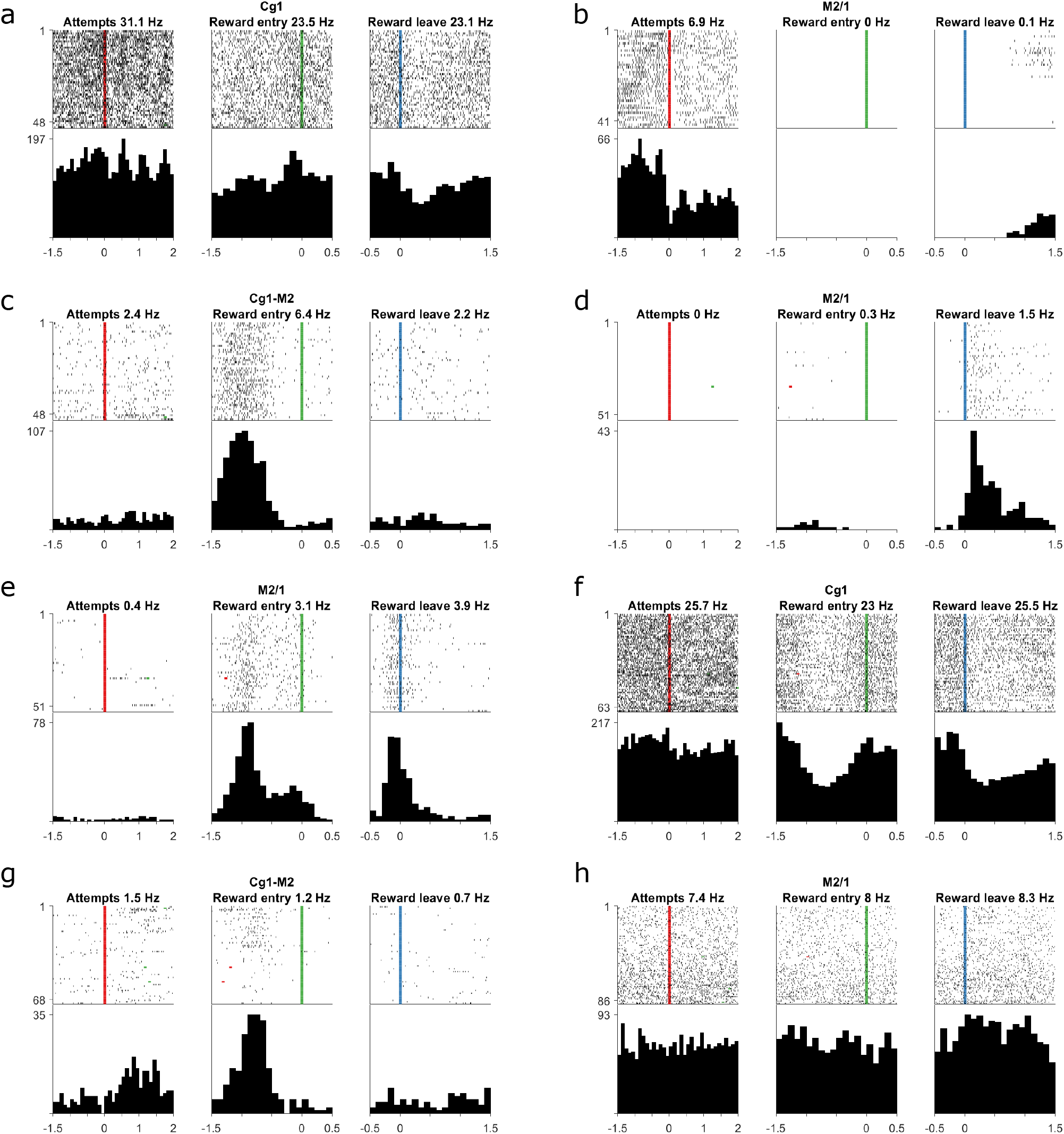
Examples of single unit activity across task-phases during PW sessions. a-h) Each panel shows the activity of one neuron across the three task-phases with a raster plot and peri-event time histogram. Neuron location (Cg1, Cg1-M2, M2/1) is indicated in the title line for each panel. For the raster plots, each row is a trial where the top-most row corresponds to the first trial and descends chronologically. Vertical lines indicate the time of event: attempt, reward entry, reward leaving. Histograms were calculated with 100 ms bins. The y-tick number indicates the highest number of spikes across all three phases. The y-axis is consistent across all three phases.

There were a total of 268 (74%) unique task-responsive neurons; 161 of these neurons were responsive to the attempt phase, 174 to reward entry, and 153 to reward leave. Ninety-seven neurons were responsive in only one epoch while 122 were responsive in two epochs and 49 were responsive in all three. There was no significant difference in the proportion of responsive neurons across the three brain locations (*X*^2^ all p’s > 0.51). Thus, unless otherwise stated, all neurons responsive to at least one task-phase were pooled together across brain locations for further analysis. Of the 92 neurons that were excluded, 56 were removed because they did not have a mean firing rate over 0.5 Hz for any task phase while the remaining 36 neurons were not responsive to any of the task phases compared to their baselines (see Methods). However, we did repeat all analyses to include the 36 non-task responsive neurons and found no qualitative differences in the results. Of the 268 neurons, 41 were classified as interneurons by their overall mean firing rate being greater than 10 Hz. The response profiles of the putative pyramidal neurons and interneurons can be seen in Supplemental Table 1 (https://doi.org/10.6084/m9.figshare.14755929.v1).

The normalized, trial averaged activity from these task-responsive neurons is shown in Figure 4a. Trials were defined as the sequence of events beginning at a rope pull attempt followed by reward zone entry and concluding with reward zone leaving (see Methods). Neural activity during the attempt phase was highly correlated across task-responsive neurons which may reflect tonic activity patterns during the rope pulling action (Figure 4b). In contrast, neural activity during reward entry and reward leaving was more phasic with lower correlation values from early compared to later time points within these phases. Activity in the attempt phase was highly dissimilar compared to the two reward phases. To further visualize neural activity across trials, we carried out PCA on the trial averaged activity of the task-responsive neurons (Figure 4c). Supporting the cross-correlation analysis, principal component 1 (PC1; 32% of variance) differentiated trial-bins by attempt phase versus the two reward phases (reward entry and reward leave). PC2 and PC3 explained a further 16% and 11% of the observed variance, respectively. Trial-bins in the attempt phase clustered together before transitioning to the reward phases. From the start of reward entry, the trajectory traces out a path from reward entry to the end of reward leaving; there is some clustering around the transition point from reward entry to reward leaving that may reflect reward consumption (see Figure 4c). Overall, dmPFC neurons track task states; ensemble activity is sustained during attempts and then smoothly transitions through the reward entry and leaving phases.

**Figure 4:**
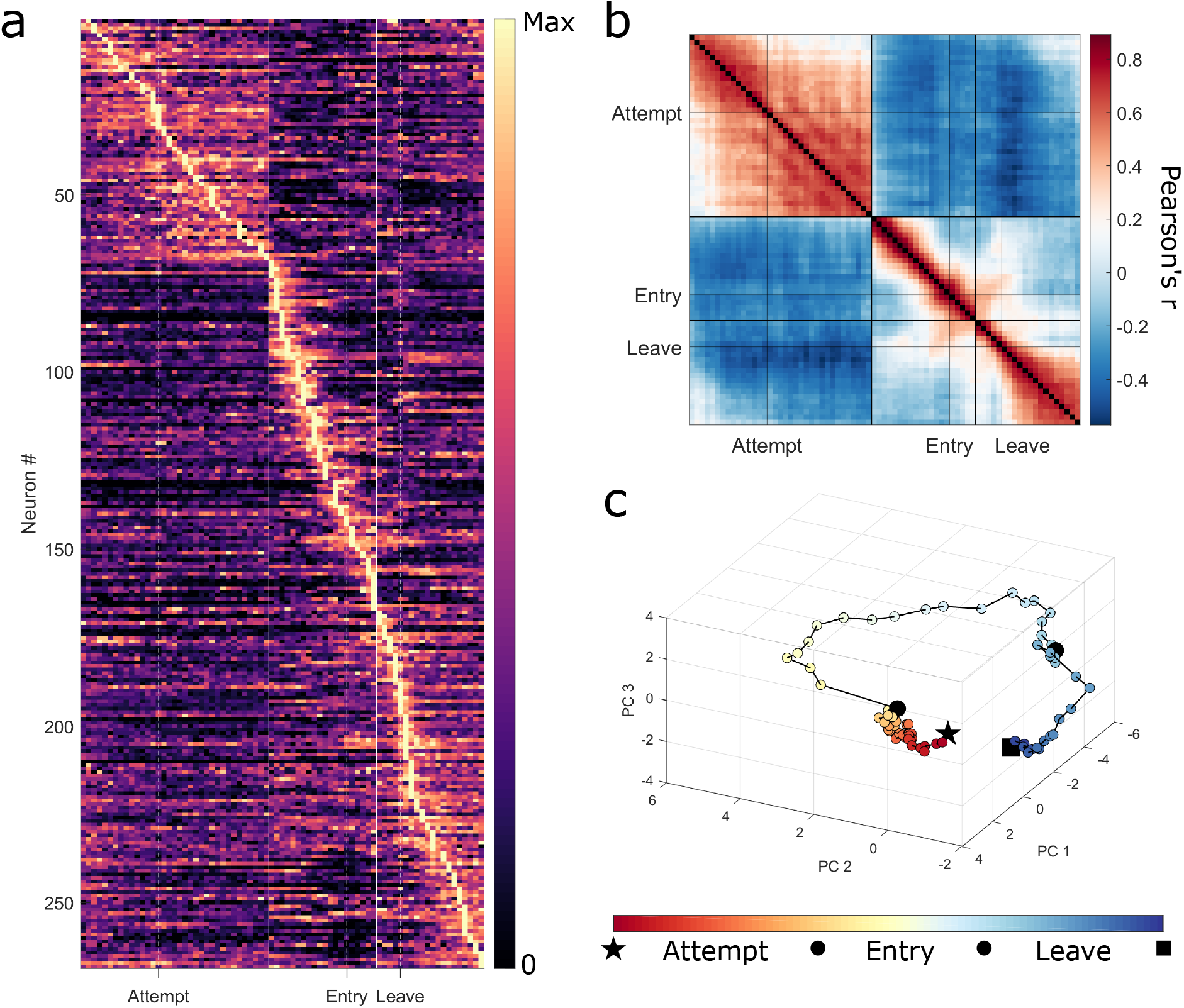
Population activity of task-responsive neurons during the PW paradigm. a) Trial averaged activity of all 268 task-responsive neurons, sorted by trial-bin in which neurons had their highest activity. Each trial was segmented into 100ms time bins. For graphical purposes, each neuron’s trial averaged activity was scaled between 0 and 1. b) Cross-correlation matrix of the average, normalized activity of all active neurons across trial phases. c) PCA scores from the average neural activity across trials in the first three principal components. Every point represents the top three PC scores from one 100ms bin. Red represents the start of the trial and progresses to blue at the end of the trial. The first time bin is highlighted with a star and last bin with a square. Black circles represent trial-phase transitions from Attempt to Reward Entry, and Reward Entry to Reward Leave.

To test whether neural activity reflected differences in future behavior, we parsed trials based on whether or not the rat went on to engage in another rope pull attempt after reward consumption. A trial was deemed an “engage” trial if – after the reward leave phase – the rat went on to perform another rope pull within 12 seconds. Twelve seconds was chosen as a cut-off since, on approximately 75% of all trials, rats made another attempt within 12 seconds (Figure 5a). The PCA trial-phase trajectories for engage versus disengage trials were highly similar (PC1, 24%; PC2, 11%; PC3, 8%; Figure 5b) and distances between bins tended to be within 2 S.D. of the mean distance between randomly sampled trajectories (Figure 5c). However, towards the end of the Reward leave phase, the engage and disengage trajectories started to diverge. There was a main effect for task-phase (F (2, 72) = 34.3, p < 0.0001). Post-hoc pairwise testing showed no significant difference between the distances of engage-disengage trajectories for attempt versus reward entry (p = 0.77) but reward leaving distances were significantly greater compared to both the attempt phase (p < 0.0001) and reward entry (p < 0.0001). Thus, in dmPFC the choice to engage or disengage from the task emerges relatively late as rats leave the reward zone.

**Figure 5:**
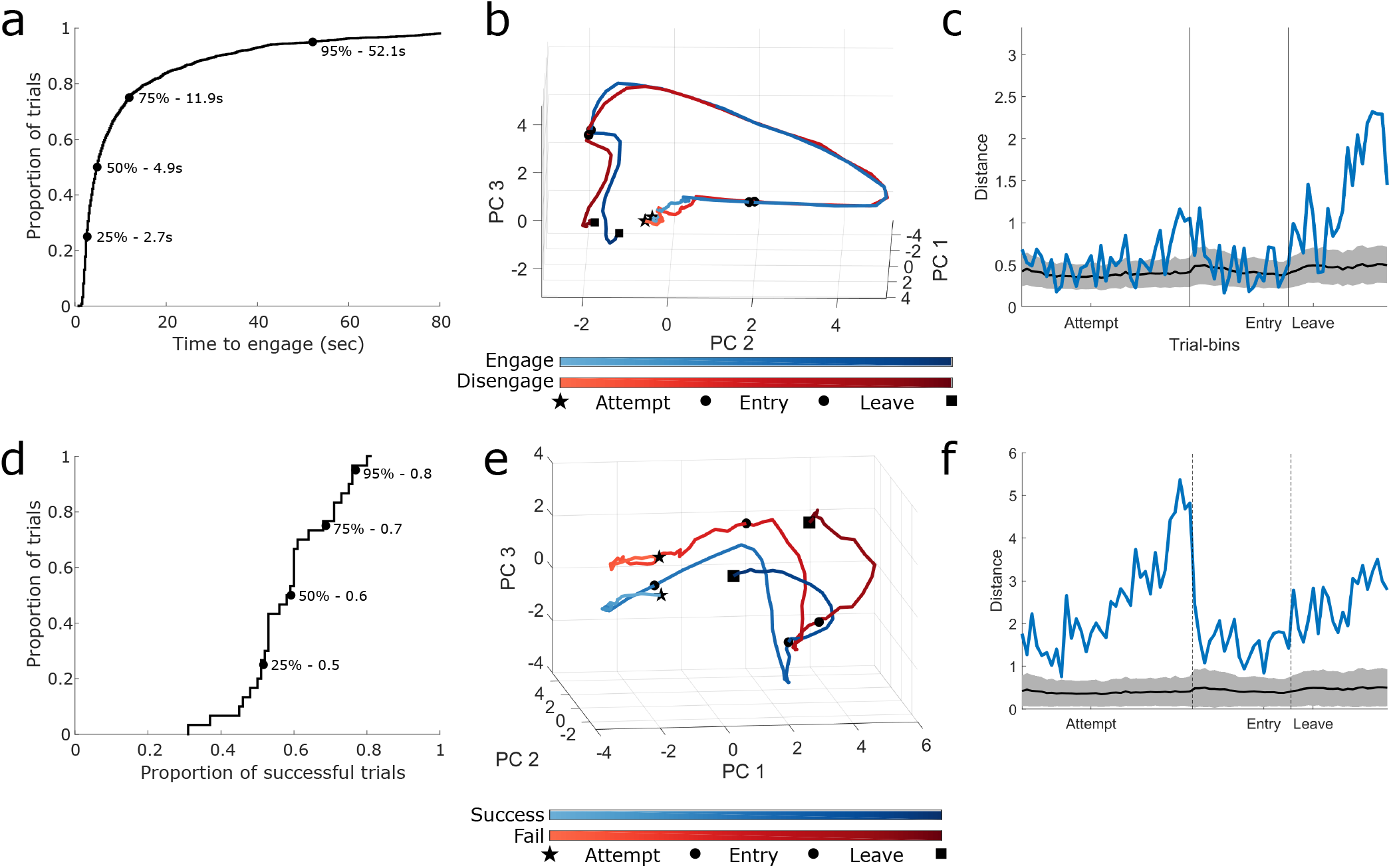
dmPFC neural activity changes during task engagement and failure in the PW paradigm. a) Cumulative distribution function of the time it took rats to engage in another trial following reward zone leaving. Quartile boundaries indicated by text. b) Neural trajectories in PC space during trials where rats subsequently engaged in a rope pulling trial (blue) or disengaged (red). c) Distances between engage vs disengage trials in PC space (blue line) for each 100ms time bin. Solid black line is the mean distance between random trajectories and grey area represents ± 2 S.D. d) Cumulative distribution function for the proportion of successful trials across sessions. Quartile boundaries indicated by text. e) Neural trajectories in PC space during trials where rats succeeded in lifting the weight 30 cm (blue) or failed (red). f) Distances between successful vs failed trials in PC space (blue line) for each 100ms time bin. Solid black line is the mean distance between random trajectories and grey area represents ± 2 S.D. b & e) Lighter color indicates earlier within the trial. The first time bin is highlighted with a star and last bin with a square. Black circles represent trial-phase transitions from Attempt to Entry and Entry to Leave.

To test whether neural activity reflected differences in rope pulling outcome, we also parsed trials based on whether or not the trial was a “success” (i.e., the weight was lifted 30 cm to trigger reward dispensing) or a “failure” (i.e., rope pulling was initiated but the weight was not lifted to the full required height of 30 cm). For 75% of recording sessions the average success rate was above 50% (Figure 5d). The PCA trial phase trajectories of successful versus failed trials were distinct but had similar overall shapes (PC1, 18%; PC2, 8%; PC3, 7%; Figure 5e). The distances between successful and failed time bins in PC space were greater than 2 S.D. of randomly sampled trajectories (Figure 5f). Task phase had a significant effect on successful versus failed trajectory distances (F (2, 72) = 8.68, p = 0.0004) where both the attempt phase (p < 0.0008) and reward leaving phase (p = 0.002) had greater distances than reward entry. Thus, dmPFC ensemble states may reflect levels of task engagement and the amount of effort put into a trial.

To test whether neural activity in each task-phase was changing over time as rats carried out the PW paradigm, we parsed trials based on their temporal quartile of occurrence. Thirty-five percent (57/161) of attempt phase neurons, 32% (56/174) of reward entry neurons, and 22% (34/153) of reward leave neurons had a significant effect for quartile. Of the neurons with a quartile effect, we determined in which quartile they had the highest firing rate. For all three task-phases, the observed frequencies of peak activity across the quartiles were significantly different than an equal distribution across the quartiles (Attempts, *X*^2^(3) = 28.7, p < 0.0001; Entry, *X*^2^(3) = 14.7, p = 0.0021; Leaving, *X*^2^(3) = 20.8, p < 0.0002; Figure 6a). The majority of neurons, across task-phases had their highest firing rates in Q1. We also analyzed each rat’s neurons individually and the majority (3/5) of rats had the highest proportion of neurons most active in Q1. To test if neural activity was changing over the quartiles, we carried out one-way repeated measures ANOVAs on the population of task-active quartile-effect neurons in each task-phase (Figure 6b). Across the four quartiles, there was a significant effect for quartile for the attempt phase (F (1.8, 103.3) = 12.4, p < 0.0001), reward entry (F (2.3, 128.3) = 3.2, p = 0.037), and reward leave phase (F (2.0, 66.6) = 10.2, p < 0.0002). Specifically, pairwise post-hoc testing showed Q1 had significantly higher neural activity compared to Q3 and Q4 during attempts and reward leaving (all p’s < 0.05), but not during reward entry (no pairwise comparisons were significant). For all three task-phases, we did not see a difference in the observed numbers of quartile selective neurons across brain regions (Chi-square test, all p’s > 0.28; Figure 6c). We tested whether the magnitude of firing rate change was impacted by session duration and found no effect of session duration for all three task phases (all p’s > 0.23, R^2^’s < 0.008; Supplemental Figure S2a, https://doi.org/10.6084/m9.figshare.14515659). These data may indicate that a subpopulation of dmPFC neurons have their peak activity early on in the session and then decrease firing over the session.

**Figure 6:**
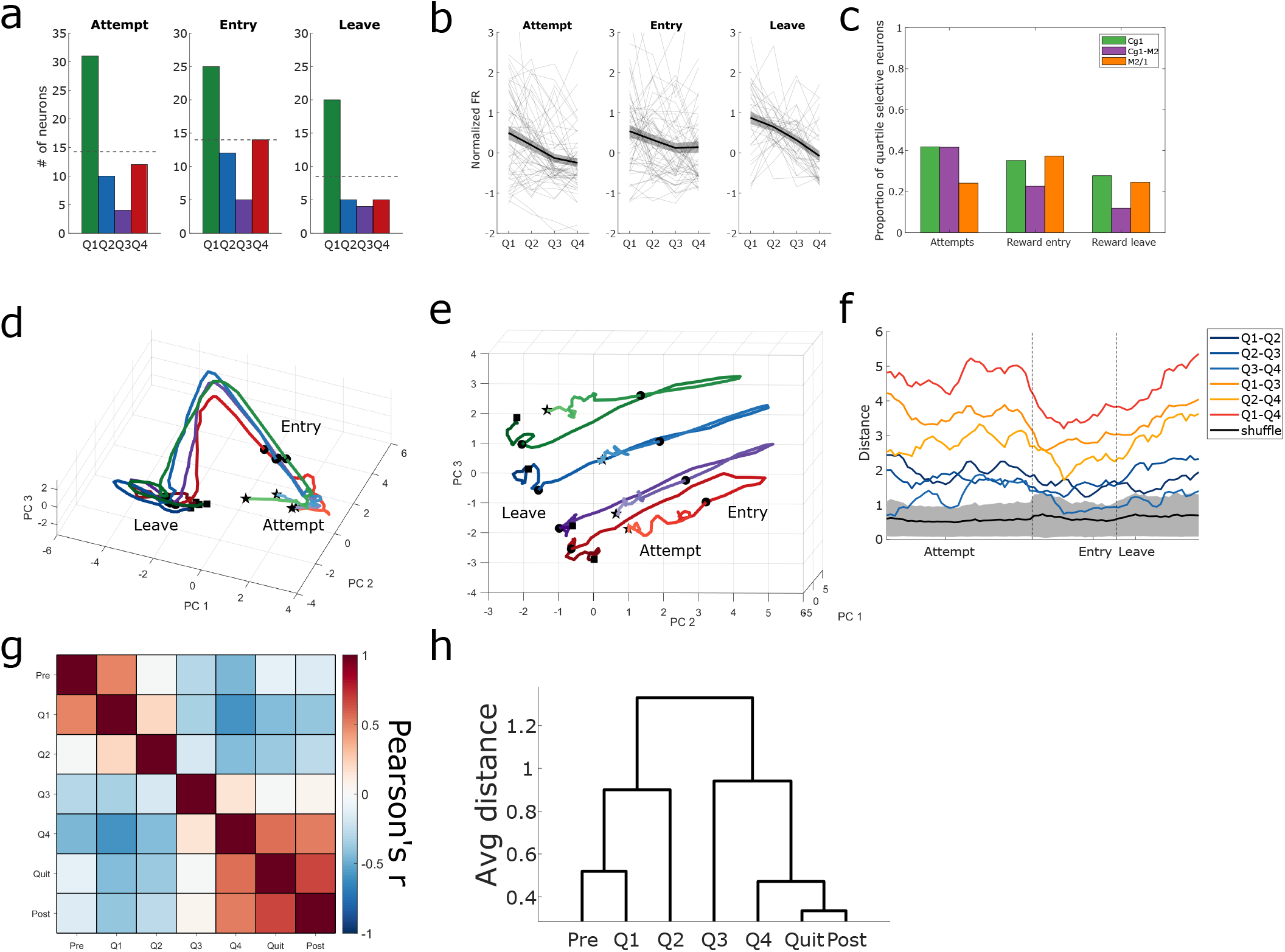
dmPFC neural activity across quartiles during the PW paradigm. a) Distributions of peak neural activity across the four quartiles for each task-phase. Dotted line indicates equal distribution across quartiles. b) Average population activity of task-active neurons with a quartile effect. Activity averaged over all neurons shown as solid black line with grey shading representing S.E.M. Individual neuron activity plotted as light grey lines. c) Proporti on of task-responsive neurons with a significant quartile effect by brain region. d & e) PCA of trial activity across the four quartiles, viewed from two different angles. Q1, green; Q2, blue; Q3, purple; Q4, red. f) Pairwise distances between PC trajectories (colored) and average distance between shuffled quartiles (black) with ± 2 S.D. in grey. Cooler colors represent temporally closer pairs while warmer colors indicate more temporally distant pairs. g) Cross-correlation matrix of 2 min epoch activity across sessions. Pre, pre-baseline; post, post-baseline. h) Dendrogram of hierarchically clustered correlation distances.

To visualize any changes in neural activity over the quartiles, we carried out PCA on the trial averaged activity of all 268 task-responsive neurons across the quartiles (PC1, 17%; PC2, 8%; PC3, 7%). The PCA trial-phase trajectories were highly conserved across quartiles (Figure 6d). Similar to the PCA trial phase trajectory that was observed when all trials across a session were averaged together (see Figure 4c), each quartile’s trajectory had activity clustered tightly during attempts, then abrupt shifts to reward entry and then a smooth transition through reward entry to reward leaving (Figure 6d). PC3, however, appears to differentiate the trajectories by quartile in that distinct trajectories for each of the four quartiles can be seen (Figure 6e, f). Pairwise distances for every trial-bin in PC space across the four quartiles showed a significant effect for quartile pairs (F (5, 444) = 370, p < 0.0001; Figure 6f). The distances between Q1-Q2 and Q2-Q3 were equivalent (p = 0.99) whereas all other pairwise comparisons were significantly different from one another (all p’s < 0.0001). Q3-Q4 had the closest trajectories, whereas Q1-Q4 were the most distant. All distances from real trajectories were significantly greater than those from randomly generated quartiles (paired t-tests, all p’s < 0.0001). The relative distances between pairs of quartiles remain even when the population of neurons is subsampled down to 10% of the original number of neurons (Supplemental Figure S3, https://doi.org/10.6084/m9.figshare.14515665.v1). We repeated the PCA for the five individual rats that had more than 30 neurons. The general trend of more-distant quartiles having more-distant trajectories was seen in 3/5 rats (data not shown). For all five rats, the Q1-Q4 comparison had the farthest distances. Thus, across sessions the dmPFC may have a consistent representation of the task-relevant behavioral states that is modified over time by the current task utility.

To determine if neural activity changes could be seen more broadly across a recording session – as rats went from entering the apparatus, to starting the task, to quitting the task – we averaged the activity of the 268 task-active neurons in 2 minute epochs. These epochs consisted of the pre-baseline period, the four quartiles of task performance, the quit period where the rat could continue to do the task (rope available) but chose not to, and the final post-baseline period (see Methods). We predicted that neural activity would be the most similar when comparing the pre-baseline, quit period, and post-baseline epochs, as behaviorally the rat is not engaged in the task during any of these periods. Cross-correlation matrices revealed two distinct patterns which were contrary to our predictions: neural activity was highly similar across the pre-baseline, Q1, and Q2 epochs, and highly similar across the Q3, Q4, quit period, and post-baseline epochs (Figure 6g). Specifically, neural activity was highly correlated between pre-baseline and Q1 (r = 0.48), Q4 and the quit period (r = 0.54), as well as Q4 and the post-baseline period (r = 0.51). Clustering analyses confirmed these patterns showing two distinct clusters relating to the first half of the session and the latter half of the session (Figure 6h). The differences in firing rate activity between Q1 and Q4 in these analyses was significantly influenced by recording duration (linear regression; F = 5.52, p = 0.0195; Supplemental Figure S2c, https://doi.org/10.6084/m9.figshare.14515659). However, the model fit was poor (R^2^ = 0.02). Thus, neural activity in the dmPFC appears to shift as the session progresses, irrespective of the specific behaviors being performed.

We repeated the above analyses on the 227 putative pyramidal neurons and the 41 putative interneurons. The task-responsiveness of pyramidal neurons and interneurons can be seen in Supplemental Table 1 (https://doi.org/10.6084/m9.figshare.14755929.v1). The pyramidal neuron restricted dataset yielded very similar results to that of the whole population and all statistical testing had the same outcomes for peak quartile across task phases, decrease in firing rate over quartiles, and task phase responsiveness by brain region (Supplemental Figure 4; https://doi.org/10.6084/m9.figshare.14757342).

In contrast, the interneuron restricted dataset did differ from the total and pyramidal populations (Supplemental Figure 5; https://doi.org/10.6084/m9.figshare.14757876.v1). Mainly, while interneurons were highly task responsive as shown by the proportion of neurons selective to the three task phases (Supplemental Table 1), the neurons did not show a change in firing rate across quartiles (all P’s > 0.2; Supplemental Figure 5b). Our other statistical tests were no longer appropriate (e.g., Chi-square tests) due to the low sample sizes (Supplemental Figure 5a, c). The lack of robust decrease in firing rate across the quartiles was also evident in the principal component analysis where PC1 (33% of variance), PC2 (11%), and PC3 (8%) reflected the three task phases (Supplemental Figure 5d, e) while PC4 (6%) showed modest quartile differences (Supplemental Figure 5f, g). Across the entire session, the interneuron population correlations were similar to the overall and pyramidal neuron data sets (Supplemental Figure 5h, i). Overall, interneurons may have a specialized role in encoding the task structure (task phases) and show less modulation by task utility than pyramidal neurons.

While there were no differences in the proportion of neurons active for a given task phase across brain regions (Figure 6c), we repeated our analyses on each brain area individually. The normalized trial activity of all the neurons from each brain regions is shown in Supplemental Figure 6a-c (https://doi.org/10.6084/m9.figshare.14758143.v2). PCA was carried out on the population of neurons from each brain region across quartiles during the three task phases (Supplemental Figure 6d-f). The general trend of the top principal components corresponding to differentiating the task phases was true for all three brain regions. However, when comparing the pairwise distances between PC trajectories across quartiles, Cg1 (Supplemental Figure 6g) and M2/1 (Supplemental Figure 6i) had more distinct trajectories for each quartile than the intermediary Cg1-M2 (Supplemental Figure 6h). However, when looking at the neural responses over large, 2-minute epochs across recording sessions, all three brain regions show the same general pattern of closer in time epochs being more similar (Supplemental Figure 6j-l). These data may suggest that there are subtle differences in the responsiveness of neurons in these different brain areas, though targeted manipulation would be necessary to determine their functional roles.

### Neuronal task selectivity – Fixed weight paradigm

The PW paradigm affords an animal continuously decreasing utility, due primarily to the increasing effort costs of the increasing weight. Rats must decide when to quit the task when the weight becomes too physically demanding. However, quitting in the PW task may be spurred by other primary factors such as fatigue or satiation. In order to try and understand the neural activity behind quitting behaviors more broadly, we analyzed dmPFC neural activity as rats carried out the FW paradigm, which lacks progressive increases in weight.Instead, the utility of the FW paradigm decreases due to changes in the internal state of the animal, e.g. fatigue, motivation, satiation.

We recorded a total of 62 dmPFC neurons from the three rats that contributed the nine FW sessions shown in Figure 3 (2-4 sessions per rat). Of these 62 neurons, a total of 53 (86%) met our task-responsive criteria for further analysis; trial averaged activity is shown in Figure 7a. Thirty-eight of the task-responsive neurons were active in the attempt phase, 41 in the reward entry phase, and 31 in the reward leave phase. Twelve of the task-responsive neurons were active in one phase, 25 in two phases, and 16 in all three. A cross-correlation matrix across all bins (Figure 7b) shows a similar pattern to that observed for the PW paradigm (see Figure 4b). Despite the relatively small number of neurons, PCA of the trial trajectory resulted in a similar low-dimensional trajectory (PC1, 40%; PC2, 16%; PC3, 10%; Figure 7c) to that seen for the PW trajectory (see Figure 4c). Activity clustered tightly during the attempt phase before transitioning through the reward entry phase with a relatively wide trajectory. Activity then clustered again from the reward entry to reward leave transition.

**Figure 7:**
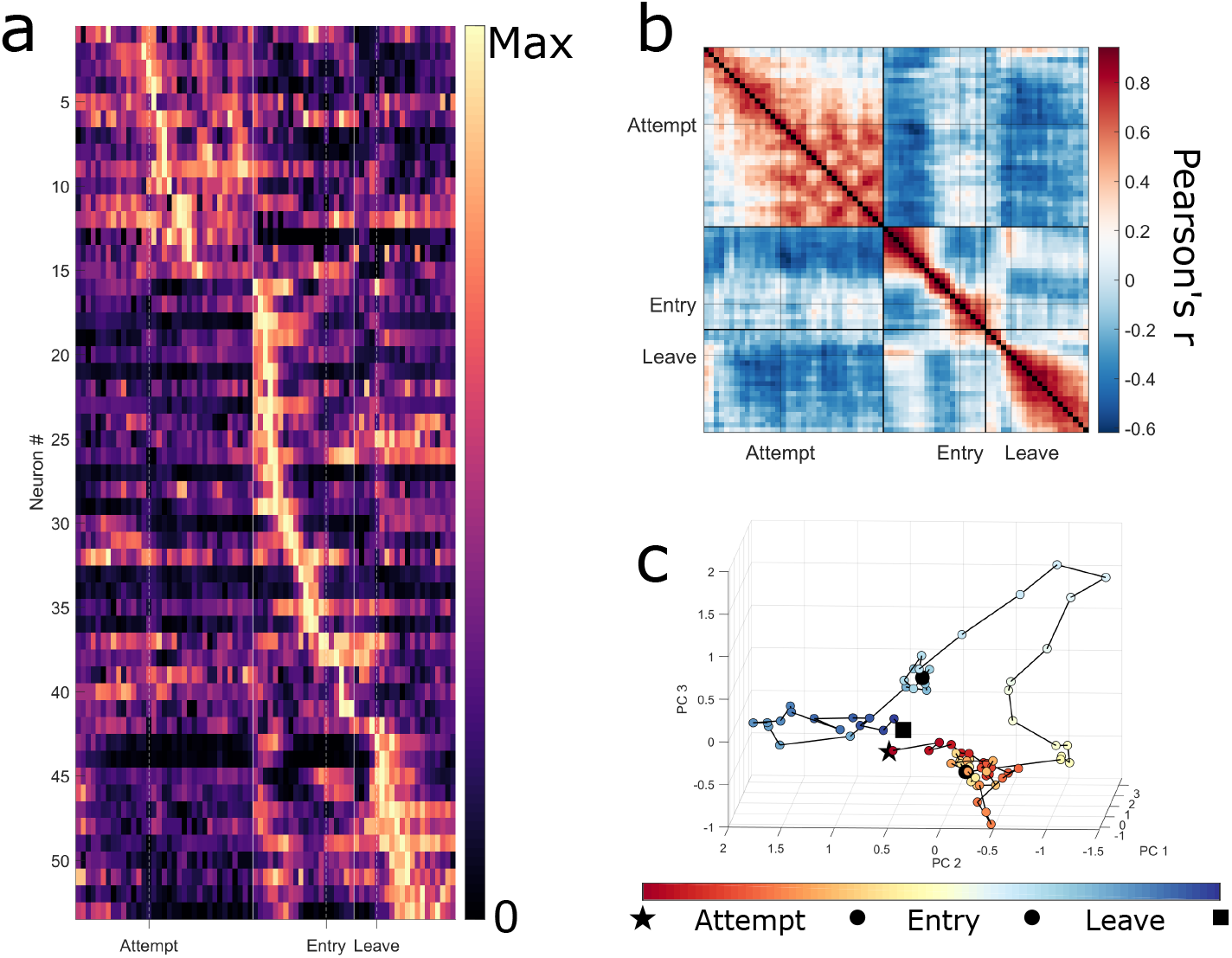
Population activity of task-responsive neurons during the FW paradigm. a) Trial averaged activity of all 53 task-responsive neurons, sorted by trial-bin in which neurons had their highest activity. Each trial was segmented into 100ms time bins. For graphical purposes, each neuron’s trial averaged activity was scaled between 0 and 1. b) Cross-correlation matrix of the average, normalized activity of all active neurons across trial phases. c) PCA scores from the average neural activity across trials in the first three principal components. Every point represents the top three PC scores from one 100ms bin. Red represents the start of the trial and progresses to blue at the end of the trial. The first time bin is highlighted with a star and last bin with a square. Black circles represent trial-phase transitions from Attempt to Reward Entry, and Reward Entry to Reward Leave.

To test whether neural activity was changing over the course of the FW paradigm, we analyzed the activity of neurons across session quartiles. Kruskal-Wallis tests were used on each task-responsive neuron to test firing rate variability across quartiles for each task-phase. Of the 38 neurons active in the attempt phase, 14 (37%) showed a significant change in firing rate across the quartiles. Twelve of the 41 (29%) reward entry and 9 of the 31 (29%) reward leave active neurons showed a significant effect for quartile. As was seen in the PW paradigm, the majority of task-active neurons with a significant effect of quartile had their highest firing rate in Q1 of the FW task (Attempts, *X*^2^(3) = 16.3, p < 0.0009; Entry, *X*^2^(3) = 10.0, p < 0.019; Leaving, *X*^2^(3) = 9.2, p < 0.027; Figure 8a). Across quartiles, the normalized firing rates from the population of task-active neurons showed a significant effect for quartile for all three task-phases (Attempts, F (2.6, 35.8) = 16.7 p = 0.001; Entry, F (2.0, 24.4) = 12.0, p = 0.005; Leaving, F (2.2, 19.6) = 12.9, p = 0.007; Figure 8b). Post-hoc pairwise comparisons revealed that the averaged population activity was significantly higher during Q1 compared to Q4 (all p’s < 0.0003) and during Q2 compared to Q4 (all p’s < 0.004) in all three task-phases. Session duration had no effect on the observed changes of firing rate from Q1 to Q4 during reward entry or leaving (p’s > 0.25). For the attempt phase, session duration was negatively correlated to the differences in firing from Q1 to Q4 (F = −2.2; p = 0.04, R^2^ = 0.11). Thus, similar to the PW paradigm, a subset of dmPFC neurons exhibit a decrease in activity across quartiles in the FW paradigm. Quartile selective neurons by brain region are shown in Figure 8c, however statistics are not provided as there were too few neurons to conduct Chi-Square tests.

**Figure 8:**
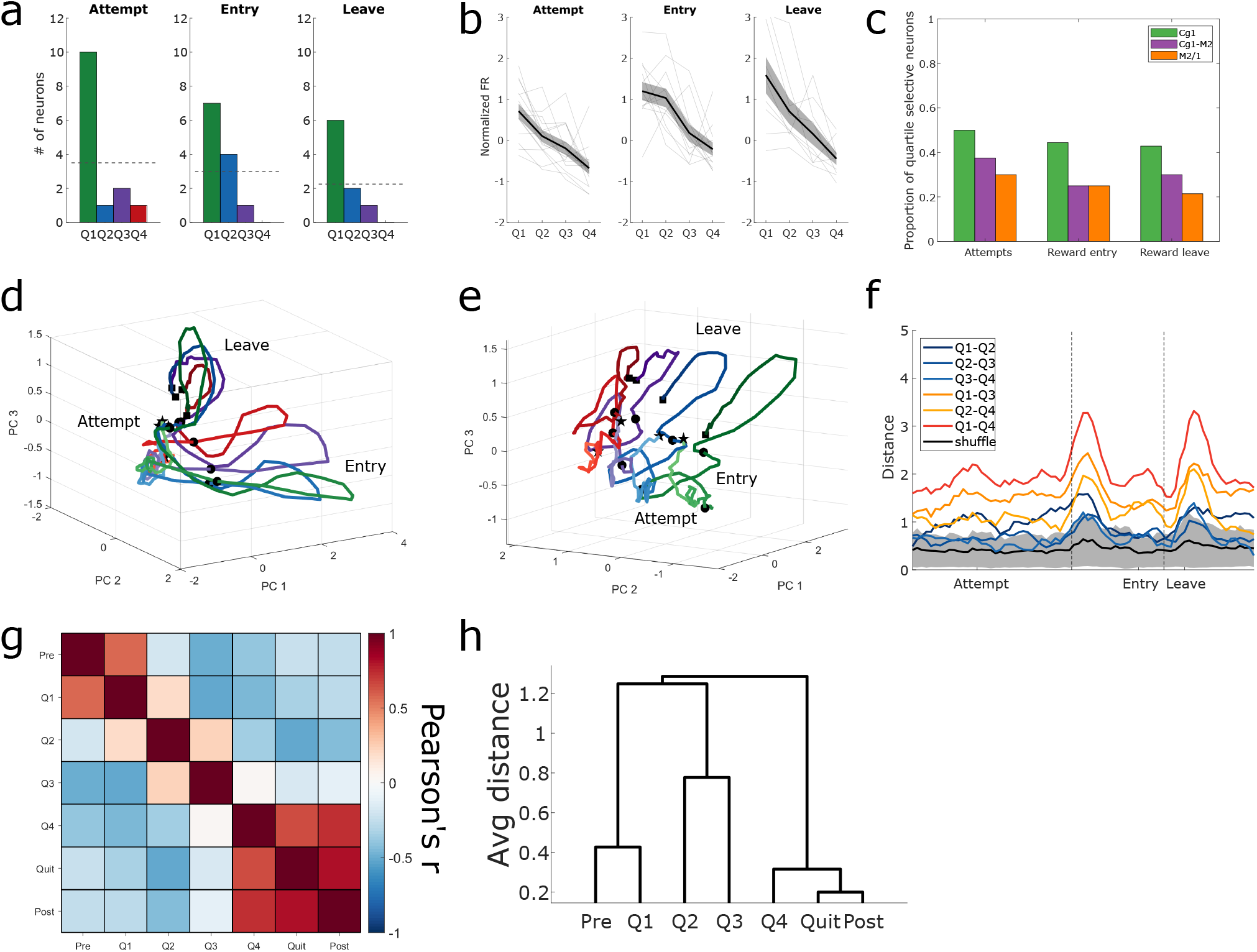
dmPFC neural activity across quartiles during the FW paradigm. a) Distributions of peak neural activity across the four quartiles for each task-phase. Dotted line indicates equal distribution across quartiles. b) Average population activity of task-active neurons with a quartile effect. Activity averaged over all neurons shown as solid black line with grey shading representing S.E.M. Individual neuron activity plotted as light grey lines. c) Proporti on of task-responsive neurons with a significant quartile effect by brain region. d & e) PCA of trial activity across the four quartiles, viewed from two different angles. Q1, green; Q2, blue; Q3, purple; Q4, red. f) Pairwise distances between PC trajectories (colored) and average distance between shuffled quartiles (black) with ± 2 S.D. in grey. Cooler colors represent temporally closer pairs while warmer colors indicate more temporally distant pairs. g) Cross-correlation matrix of 2 minute epoch activity across sessions. Pre, pre-baseline; post, post-baseline. h) Dendrogram of hierarchically clustered correlation distances.

PCA was applied to the averaged normalized activity of the 53 task-active neurons across the four quartiles during the FW task (PC1, 23%; PC2, 10%; PC3, 8%; Figure 8d, e). Generally, the same trajectory shape across the whole trial can be seen across all four quartiles (Figure 8d). However, as was seen in the PW task, the trial trajectories for each quartile can be distinguished (Figure 8e). The attempt phase remains relatively similar across quartiles, however, the wide neural trajectories during the reward entry phase show differing paths. During reward leaving, trajectories become smaller over the quartiles reflective of more stable, stationary neural states. Pairwise distances were calculated between every quartile across trial bins (Figure 8f). As was seen in the PW, temporally distant trajectories tended to be farther apart than temporally proximal trajectories (F (5, 444) = 120, p < 0.0001). Differing from the PW paradigm, distances between Q2-Q3 and Q3-Q4 (p = 0.99) and Q1-Q2 and Q2-Q4 (p = 0.15) were not different in the FW paradigm, otherwise, all other pairwise comparisons were significantly different (all p’s < 0.0001). In order of mean distance, Q1-Q2 was more distant than Q2-Q3 and Q3-Q4. Unique to the FW, distances between trajectories tended to be farthest apart prior to the reward entry point and after reward leaving point. The average distances of every real trajectory were significantly greater than those from randomly sampled trajectories (paired t-tests, all p’s < 0.0001).

Neural activity during the FW paradigm was also analyzed in broader two minute epochs across the session, which included the pre- and post-baseline periods (Figure 8g, h). Neural activity was highly correlated between pre-baseline and Q1 (r = 0.57), before becoming less correlated quartile to quartile. Neural activity across the Q4, Quit, and post-baseline periods was also highly correlated, with significant Pearson’s r values observed between Q4 and Quit (r = 0.64), Q4 and post-baseline (r = 0.72), and Quit versus post-baseline (r = 0.80). Session duration had no impact on the firing rate differences between Q1 and Q4 (F = 0.96, p = 0.34, R^2^ = 0.02). Overall, the FW paradigm results are similar to the PW results in that a consistent representation of the behavioral states of the task is modified across quartiles; in the FW paradigm, however, this cannot be directly attributed to increases in absolute physical effort across time (i.e., increments in rope weight).

## Discussion

We utilized two complimentary effortful behavioral paradigms to investigate how dmPFC neural activity responded to changes in task utility, and to test whether dmPFC ensemble activity reflected a rat’s progression towards task quitting. In both the PW and FW paradigms, behavioral metrics indicate that rats were highly engaged in the task until the final quartile (Q4) where engagement waned. In the PW paradigm, rats took longer to earn rewards and failed rope-pulling more often quartile-by-quartile as they approached their quit weight in Q4. In the FW paradigm, reward intervals and failed attempts only increased in Q4. Overall, these data show that the PW paradigm ramps up in difficulty with commensurate changes in behavior across quartiles, leading towards the quit point. In contrast, the FW paradigm presents a repetitive, consistent task where behavior is relatively stable until a change in Q4, immediately preceding the quit point.

Despite the behavioral differences between the two paradigms, we observed similar neural activity patterns. Neural activity in dmPFC neurons on average was highest in Q1 and decreased across quartiles, leading to the quit point. Since these patterns were observed in both paradigms they cannot be solely attributed to progressive increases in rope weight. PCA showed conserved neural trajectories for trial phases (attempt, reward entry, reward leave), which remained relatively consistent across the session. However, each quartile had a distinct trajectory. The more temporally close quartiles had closer PCA trajectories, with Q1 and Q4 being the most distinct in both task paradigms. Broader temporal analysis of neural activity encompassing the entire recording session – which included the baseline periods where the rope was unavailable – revealed distinct shifts in activity from the start of the session to the end. Overall, our results suggest that the dmPFC maintains a consistent representation of the sequential behavioral states of the task that is responsive to repetitive task engagement. We propose that the observed decrease in activity across a session reflects the decrease in task utility across time, and that this corresponds to waning motivation and eventual quitting of the task.

We observed similar activity patterns in neurons from Cg1, Cg1-M2, and M2/1 during WLT performance. The proportion of neurons from each brain area that were active in each task-phase were not significantly different. These results were somewhat surprising as we expected M2 neurons to be more responsive to attempt phase rope pulling and Cg1 neurons to be more responsive to reward phases. However, our results do align with the overlapping roles ascribed to the two brain areas. Generally, M2 has been proposed to select and adapt action plans based on sensory feedback (13, 15, 26) while the Cg1 monitors behaviors to determine their utility and bias behavior to maximize value (27, 28). Both brain regions may play complimentary roles in physical effort-based cost-benefit decision making: M2 selects actions and evaluates sensorimotor feedback of the costs and outcomes of those actions, while anterior Cg1 evaluates the overall utility of the selected action plan in comparison to available alternatives (18, 29). We did find some subtle differences in how strongly the regions differentiated their activity across the quartiles (Supplemental Figure 6). Further manipulation experiments, specifically spanning the medial to lateral extent of the dmPFC, would be needed to determine precise functional differences. Our results add further support that anterior Cg1 and M2 play complimentary roles in cost-benefit decision making, specifically at the intersection of action selection, action-outcome monitoring, and adjusting future behavior.

Numerous studies have shown that the Cg1/ACC and M2 (or supplementary motor area in primate) respond to the relative value of behaviors, with firing rates or BOLD responses corresponding to high value options (18, 20, 21, 23, 30–34). However, in most previous studies subjects had to choose between, or are at least were presented with, multiple stimuli or options in a given task; quitting not being one of the options (though see 7)). Furthermore, there has generally been an emphasis on the evaluation of the *relative* value of behaviors in the context of all possible behaviors (21, 31, 32, 35) even if there is no overt choice between multiple options (20, 34).

In our WLT rats could only choose between two options – carrying out the task or not; there were no other extrinsically rewarded actions available in the environment. In accordance with dmPFC encoding the relative value of a behavior, we observed a value signal that decreased over the course of the session, culminating in the quit point (quitting being the only alternative course of action). Overall our results further support the idea that dmPFC encodes the relative value of behaviors in order to choose actions that maximize utility (36) possibly through exerting cortex-wide control in task engagment (3). At a certain point, in response to changing environmental demands and internal state, quitting becomes the behavior with optimal utility when trying to minimize energy expenditure while trying to satisfy internal drives (37–40).

When we compared trials where the animal went on to engage in another rope pull, versus trials where the animal disengaged from the task, dmPFC activity in PC space showed general overlap in neural trajectories. However, significant differences were observed between engage and disengage trials after the rat left the reward cap. In separate analyses, when we compared neural activity during successful trials versus failed, non-rewarded trials, dmPFC activity in PC space showed highly distinct neural trajectories between trial types. Taken together, one interpretation of these data is that dmPFC ensemble activity drives the level of WLT engagement: one state correlates to effortful exertion and persistence in the task (successful rope-pulling and/or re-engagement in a subsequent trial), whereas a state shift correlates to faltering task performance (failed rope-pulling and/or disengagement in a subsequent trial).

Previous studies that have examined quitting tended to use progressive ratio lever pressing paradigms (41–44). Lesions to the Cg1 (43) or more ventral prelimbic cortex (44) do not alter the quit-point/breakpoint of rats in these paradigms. However, lesions to the Cg1 of rats has consistently resulted in avoidance of high-effort options (climbable-barriers, high-ratio lever presses) when relatively lower effort options are available (45–48). We have shown previously that electrical stimulation to Cg1 biases rats to quit earlier in our WLT (49).These results seem to be at odds with one another: lesioning Cg1 results in rats shifting their choice preference towards lower effort options, but does not affect high effort performance on progressive ratio tasks. Wang et al. (50) provided some clarity on this discrepancy by showing that inactivation of rat Cg1 impairs performance in a novel “do more get more” task, suggesting that Cg1 is critical for *self-paced* effort expenditure. Thus, in the context of quitting behaviors, Cg1 may provide a relative value signal of the current effortful behavior and this signal influences the decision to persist or quit in a self-paced task. Future research manipulating both Cg1 and M2 will be needed to elucidate the functional roles of the two regions in effortful decision-making tasks. Furthermore, previous evidence has shown neuromodulators such as dopamine (22, 50) and serotonin (51) influence decisions to persist or quit behaviors.

Two proposed, complimentary functions of the dmPFC are reinforcement-learning and action selection (27, 52, 53). These two functions work synergistically where the dmPFC learns about and updates a mental model of the world forming an abstract task space in order to select actions that maximize the value of an environment (54). Using PCA we demonstrated that the dmPFC contains a consistent task-space representation of behaviors during our task: effortful rope pulling (attempt phase), reward approach and consumption (reward entry phase), and prospective task engagement at the end of a trial (reward leave phase). Similar results have been reported showing the Cg1/ACC representing task-space (12, 20) including the Cg1 recalling the locations of unchosen rewards when mistakes were made, providing evidence of an addressable task-space representations (55). In addition, Wang et al. (56) reported Cg1 representations of the sequential steps of a delay-based cost-benefit task where rats were free to nose-poke for longer durations to earn proportionally larger rewards. We found similar results to Wang et al. (56) in the proportion of recorded neurons representing task states (approximately 70%) and subpopulation responding to cost-benefit contingencies or quartiles (approximately 35%). Thus, the dmPFC may have flexible, sequential representations of all task-relevant behaviors that can be modified based on cost-benefit contingencies. Together this enables optimal action selection, which sometimes means quitting the task at hand and moving on.

## References

1. Hayden BY. Economic choice: the foraging perspective. Curr Opin Behav Sci 24: 1–6, 2018. doi: 10.1016/j.cobeha.2017.12.002.

2. Stephens DW, Krebs JR. Foraging Theory. Princeton (USA): Princeton University Press, 1986.

3. Allen WE, Kauvar IV, Chen MZ, Richman EB, Yang SJ, Chan K, Gradinaru V, Deverman BE, Luo L, Deisseroth K. Global Representations of Goal-Directed Behavior in Distinct Cell Types of Mouse Neocortex. Neuron 94: 891–907.e6, 2017. doi: 10.1016/j.neuron.2017.04.017.

4. Euston DR, Gruber AJ, McNaughton BL. The Role of Medial Prefrontal Cortex in Memory and Decision Making. Neuron 76: 1057–1070, 2012. doi: 10.1016/j.neuron.2012.12.002.

5. Kennerley SW, Walton ME. Decision making and reward in frontal cortex: Complementary evidence from neurophysiological and neuropsychological studies. Behav Neurosci 125: 297– 317, 2011. doi: 10.1037/a0023575.

6. Zhao W, Kendrick KM, Chen F, Li H, Feng T. Neural circuitry involved in quitting after repeated failures: role of the cingulate and temporal parietal junction. Sci Rep 6: 24713, 2016. doi: 10.1038/srep24713.

7. Hosokawa T, Kennerley SW, Sloan J, Wallis JD. Single-Neuron Mechanisms Underlying Cost-Benefit Analysis in Frontal Cortex. J Neurosci 33: 17385–17397, 2013. doi: 10.1523/JNEUROSCI.2221-13.2013.

8. Kennerley SW, Walton ME, Behrens TEJ, Buckley MJ, Rushworth MFS. Optimal decision making and the anterior cingulate cortex. Nat Neurosci 9: 940–947, 2006. doi: 10.1038/nn1724.

9. Klein-Flügge MC, Kennerley SW, Friston K, Bestmann S. Neural signatures of value comparison in human cingulate cortex during decisions requiring an effort-reward trade-off. Neuroscience.

10. Walton ME, Bannerman DM, Alterescu K, Rushworth MFS. Functional Specialization within Medial Frontal Cortex of the Anterior Cingulate for Evaluating Effort-Related Decisions. J Neurosci 23: 6475–6479, 2003. doi: 10.1523/JNEUROSCI.23-16-06475.2003.

11. Gu X, Staines WA, Fortier PA. Quantitative analyses of neurons projecting to primary motor cortex zones controlling limb movements in the rat. Brain Res 835: 175–187, 1999. doi: 10.1016/S0006-8993(99)01576-0.

12. Lapish CC, Durstewitz D, Chandler LJ, Seamans JK. Successful choice behavior is associated with distinct and coherent network states in anterior cingulate cortex. Proc Natl Acad Sci 105: 11963–11968, 2008. doi: 10.1073/pnas.0804045105.

13. Reep RL, Corwin JV, Hashimotos A, Watsonp RT. Efferent Connections of the Rostral Portion of Medial Agranular Cortex in Rats. Brain Res Bull 19: 203–221, 1987.

14. Rolls ET. The cingulate cortex and limbic systems for emotion, action, and memory. Brain Struct Funct 224: 3001–3018, 2019. doi: 10.1007/s00429-019-01945-2.

15. Barthas F, Kwan AC. Secondary Motor Cortex: Where ‘Sensory’ Meets ‘Motor’ in the Rodent Frontal Cortex. Trends Neurosci 40: 181–193, 2017. doi: 10.1016/j.tins.2016.11.006.

16. Ebbesen CL, Insanally MN, Kopec CD, Murakami M, Saiki A, Erlich JC. More than Just a “Motor”: Recent Surprises from the Frontal Cortex. J Neurosci 38: 9402–9413, 2018. doi: 10.1523/JNEUROSCI.1671-18.2018.

17. Murakami M, Vicente MI, Costa GM, Mainen ZF. Neural antecedents of self-initiated actions in secondary motor cortex. Nat Neurosci 17: 1574–1582, 2014. doi: 10.1038/nn.3826.

18. Sul JH, Jo S, Lee D, Jung MW. Role of rodent secondary motor cortex in value-based action selection. Nat Neurosci 14: 1202–1208, 2011. doi: 10.1038/nn.2881.

19. Porter B, Hillman KL. A Novel Weight Lifting Task for Investigating Effort and Persistence in Rats. Front Behav Neurosci 13: 275, 2019. doi: 10.3389/fnbeh.2019.00275.

20. Cowen SL, Davis GA, Nitz DA. Anterior cingulate neurons in the rat map anticipated effort and reward to their associated action sequences. J Neurophysiol 107: 2393–2407, 2012. doi: 10.1152/jn.01012.2011.

21. Hillman KL, Bilkey DK. Neurons in the Rat Anterior Cingulate Cortex Dynamically Encode Cost-Benefit in a Spatial Decision-Making Task. J Neurosci 30: 7705–7713, 2010. doi: 10.1523/JNEUROSCI.1273-10.2010.

22. Salamone JD, Cousins MS, Bucher S. Anhedonia or anergia? Effects of haloperidol and nucleus accumbens dopamine depletion on instrumental response selection in a T-maze cost/benefit procedure. Behav Brain Res 65: 221–229, 1994. doi: 10.1016/0166-4328(94)90108-2.

23. Kennerley SW, Wallis JD. Evaluating choices by single neurons in the frontal lobe: outcome value encoded across multiple decision variables. Eur J Neurosci 29: 2061–2073, 2009. doi: 10.1111/j.1460-9568.2009.06743.x.

24. Hyman JM, Ma L, Balaguer-Ballester E, Durstewitz D, Seamans JK. Contextual encoding by ensembles of medial prefrontal cortex neurons. Proc Natl Acad Sci 109: 5086–5091, 2012. doi: 10.1073/pnas.1114415109.

25. Paxinos G, Watson C. The rat brain in stereotaxic coordinates. Amsterdam; Boston: Academic Press/Elsevier, 2007.

26. Olson JM, Li JK, Montgomery SE, Nitz DA. Secondary Motor Cortex Transforms Spatial Information into Planned Action during Navigation. Curr Biol 30: 1845–1854.e4, 2020. doi: 10.1016/j.cub.2020.03.016.

27. Holroyd CB, McClure SM. Hierarchical control over effortful behavior by rodent medial frontal cortex: A computational model. Psychol Rev 122: 54–83, 2015. doi: 10.1037/a0038339.

28. Shenhav A, Botvinick MM, Cohen JD. The Expected Value of Control: An Integrative Theory of Anterior Cingulate Cortex Function. Neuron 79: 217–240, 2013. doi: 10.1016/j.neuron.2013.07.007.

29. Sul JH, Kim H, Huh N, Lee D, Jung MW. Distinct Roles of Rodent Orbitofrontal and Medial Prefrontal Cortex in Decision Making. Neuron 66: 449–460, 2010. doi: 10.1016/j.neuron.2010.03.033.

30. Croxson PL, Walton ME, O’Reilly JX, Behrens TEJ, Rushworth MFS. Effort-Based Cost-Benefit Valuation and the Human Brain. J Neurosci 29: 4531–4541, 2009. doi: 10.1523/JNEUROSCI.4515-08.2009.

31. Hart EE, Blair GJ, O’Dell TJ, Blair HT, Izquierdo A. Chemogenetic Modulation and Single-Photon Calcium Imaging in Anterior Cingulate Cortex Reveal a Mechanism for Effort-Based Decisions. J Neurosci 40: 5628–5643, 2020. doi: 10.1523/JNEUROSCI.2548-19.2020.

32. Hillman KL, Bilkey DK. Neural encoding of competitive effort in the anterior cingulate cortex. Nat Neurosci 15: 1290–1297, 2012. doi: 10.1038/nn.3187.

33. Kennerley SW, Dahmubed AF, Lara AH, Wallis JD. Neurons in the Frontal Lobe Encode the Value of Multiple Decision Variables. J Cogn Neurosci 21: 1162–1178, 2009. doi: 10.1162/jocn.2009.21100.

34. Porter BS, Hillman KL, Bilkey DK. Anterior cingulate cortex encoding of effortful behavior. J Neurophysiol 121: 701–714, 2019. doi: 10.1152/jn.00654.2018.

35. Wallis JD, Kennerley SW. Contrasting reward signals in the orbitofrontal cortex and anterior cingulate cortex: Contrasting reward signals in the orbitofrontal cortex. Ann N Y Acad Sci 1239: 33–42, 2011. doi: 10.1111/j.1749-6632.2011.06277.x.

36. Rushworth M, Walton M, Kennerley S, Bannerman D. Action sets and decisions in the medial frontal cortex. Trends Cogn Sci 8: 410–417, 2004. doi: 10.1016/j.tics.2004.07.009.

37. Allen WE, Chen MZ, Pichamoorthy N, Tien RH, Pachitariu M, Luo L, Deisseroth K. Thirst regulates motivated behavior through modulation of brainwide neural population dynamics. Science 364: 11, 2019.

38. Hockey R. The psychology of fatigue: work, effort, and control. Cambridge: Cambridge University Press, 2013.

39. Noakes TD. Fatigue is a Brain-Derived Emotion that Regulates the Exercise Behavior to Ensure the Protection of Whole Body Homeostasis. Front Physiol 3, 2012. doi: 10.3389/fphys.2012.00082.

40. van der Linden D. The urge to stop: The cognitive and biological nature of acute mental fatigue. In: Cognitive fatigue: Multidisciplinary perspectives on current research and future applications., edited by Ackerman PL. American Psychological Association, p. 149–164.

41. Hodos W. Progressive Ratio as a Measure of Reward Strength. Sci New Ser 134: 943–944, 1961.

42. Rickard JF, Body S, Zhang Z, Bradshaw CM, Szabadi E. Effect of reinforecer magnitude on performance maintained by progressive-ratio schedules. J Exp Anal Behav 91: 75–87, 2009. doi: 10.1901/jeab.2009.91-75.

43. Schweimer J. Involvement of the rat anterior cingulate cortex in control of instrumental responses guided by reward expectancy. Learn Mem 12: 334–342, 2005. doi: 10.1101/lm.90605.

44. Swanson K, Goldbach HC, Laubach M. The rat medial frontal cortex controls pace, but not breakpoint, in a progressive ratio licking task. Behav Neurosci 133: 385–397, 2019. doi: 10.1037/bne0000322.

45. Hart EE, Gerson JO, Zoken Y, Garcia M, Izquierdo A. Anterior cingulate cortex supports effort allocation towards a qualitatively preferred option. Eur J Neurosci 46: 1682–1688, 2017. doi: 10.1111/ejn.13608.

46. Hauber W, Sommer S. Prefrontostriatal Circuitry Regulates Effort-Related Decision Making. Cereb Cortex 19: 2240–2247, 2009. doi: 10.1093/cercor/bhn241.

47. Holec V, Pirot HL, Euston DR. Not all effort is equal: the role of the anterior cingulate cortex in different forms of effort-reward decisions. Front Behav Neurosci 8, 2014. doi: 10.3389/fnbeh.2014.00012.

48. Rudebeck PH, Walton ME, Smyth AN, Bannerman DM, Rushworth MFS. Separate neural pathways process different decision costs. Nat Neurosci 9: 1161–1168, 2006. doi: 10.1038/nn1756.

49. Silva C, Porter BS, Hillman KL. Stimulation in the Rat Anterior Insula and Anterior Cingulate During an Effortful Weightlifting Task. Front Neurosci 15: 643384, 2021. doi: 10.3389/fnins.2021.643384.

50. Wang S, Hu S-H, Shi Y, Li B-M. The roles of the anterior cingulate cortex and its dopamine receptors in self-paced cost–benefit decision making in rats. Learn Behav 45: 89–99, 2017. doi: 10.3758/s13420-016-0243-0.

51. Lottem E, Banerjee D, Vertechi P, Sarra D, Lohuis M oude, Mainen ZF. Activation of serotonin neurons promotes active persistence in a probabilistic foraging task. Nat Commun 9: 1000, 2018. doi: 10.1038/s41467-018-03438-y.

52. Hadland KA, Rushworth MFS, Gaffan D, Passingham RE. The Anterior Cingulate and Reward-Guided Selection of Actions. J Neurophysiol 89: 1161–1164, 2003. doi: 10.1152/jn.00634.2002.

53. Paus T. Primate anterior cingulate cortex: Where motor control, drive and cognition interface. Nat Rev Neurosci 2: 417–424, 2001. doi: 10.1038/35077500.

54. Heilbronner SR, Hayden BY. Dorsal Anterior Cingulate Cortex: A Bottom-Up View. Annu Rev Neurosci 39: 149–170, 2016. doi: 10.1146/annurev-neuro-070815-013952.

55. Mashhoori A, Hashemnia S, McNaughton BL, Euston DR, Gruber AJ. Rat anterior cingulate cortex recalls features of remote reward locations after disfavoured reinforcements. eLife 7: e29793, 2018. doi: 10.7554/eLife.29793.

56. Wang S, Shi Y, Li B-M. Neural representation of cost–benefit selections in rat anterior cingulate cortex in self-paced decision making. Neurobiol Learn Mem 139: 1–10, 2017. doi: 10.1016/j.nlm.2016.12.003.

